# Gut microbial structural variations as determinants of human bile acid metabolism

**DOI:** 10.1101/2021.02.28.432952

**Authors:** Daoming Wang, Marwah Doestzada, Lianmin Chen, Sergio Andreu-Sánchez, Inge C.L. van den Munckhof, Hannah Augustijn, Martijn Koehorst, Vincent W. Bloks, Niels P. Riksen, Joost H.W. Rutten, Mihai G. Netea, Alexandra Zhernakova, Folkert Kuipers, Jingyuan Fu

## Abstract

Bile acids (BAs) facilitate intestinal fat absorption and act as important signaling molecules in host□gut microbiota crosstalk. BA-metabolizing pathways in the microbial community have been identified, but how the highly variable genomes of gut bacteria interact with host BA metabolism remains largely unknown. We characterized 8,282 structural variants (SVs) of 55 bacterial species in the gut microbiomes of 1,437 individuals from two Dutch cohorts and performed a systematic association study with 39 plasma BA parameters. Both variations in SV-based continuous genetic makeup and discrete subspecies showed correlations with BA metabolism. Metagenome-wide association analysis identified 797 replicable associations between bacterial SVs and BAs and SV regulators that mediate the effects of lifestyle factors on BA metabolism. This is the first large-scale microbial genetic association analysis to demonstrate the impact of bacterial SVs on human BA composition, and highlights the potential of targeting gut microbiota to regulate BA metabolism through lifestyle intervention.

## Introduction

Bile acids (BAs) represent an important class of biologically-active metabolites that act at the interface between host and gut microbiota. BAs are amphiphilic steroids synthesized from cholesterol in the liver and are well-known for their roles in facilitating intestinal fat absorption, promoting hepatic bile formation and maintaining whole-body cholesterol balance. In addition, BAs exert hormone-like functions by signaling via membrane-bound and nuclear receptors involved in the control of lipid, glucose and energy metabolism (Kuipers et al., 2014). Altered BA metabolism has been associated with several metabolic diseases, including type 2 diabetes (T2D) and non-alcoholic fatty liver disease (NAFLD) (Chávez-Talavera et al., 2017), and with colorectal cancer (Dermadi et al., 2017) and hepatocellular carcinoma (Gao et al., 2019).

Gut bacteria are essential players in human BA metabolism: bacterial bile salt hydrolases (BSH) convert the glycine- and taurine-conjugated primary BAs produced by the liver (cholic (CA) and chenodeoxycholic (CDCA) acids) into unconjugated primary BAs that can subsequently be dehydroxylated to form secondary BAs (deoxycholic (DCA) and lithocholic (LCA) acids) (Jia et al., 2017). Secondary BAs are efficiently absorbed in the ileum and, to a lesser extent the colon, and return to the liver via the portal venous system for re-secretion into the bile. Consequently, the BA pool consists of a mixture of primary and secondary BAs that travel between liver and intestine within the enterohepatic circulation. Recent cohort studies have demonstrated remarkable inter-individual variation in the human BA pool composition, as inferred by analyzing BA composition in peripheral blood from both healthy (Steiner et al., 2011) and obese subjects (Chen et al., 2020). Importantly, this variability could largely be attributed to metabolic and transport processes within the enterohepatic circulation rather than to differences in hepatic synthesis rates, implying a key role for the microbiome in BA diversity (Chen et al., 2020).

In recent years, we have learned a great deal about the large variability of microbial composition in healthy humans and compositional alterations associated with specific diseases (Falony et al., 2016; Jackson et al., 2018; Zhernakova et al., 2016). The relationships between host BA pool and gut microbial composition were also investigated in several cohorts with differing health status. For instance, in untreated T2D patients, concentrations of plasma glycoursodeoxycholic acid (GUDCA), LCA and DCA were associated with the overall composition of gut microbiome (Gu et al., 2017). In the 300OB obesity cohort, a considerable number of associations were identified between the relative abundances of gut bacteria and BA parameters in feces and plasma (Chen et al., 2020). Many bacterial genes involved in BA biotransformation have been identified through experimental and homologue-based bioinformatic approaches. For instance, the BSH gene pool has been quantified and characterized in the metagenomes of diverse populations (Song et al., 2019). Based on the catalog of known BA-related genes present in gut bacterial genomes, the BA biotransformation potential of individuals can be predicted using metabolic model reconstructions (Heinken et al., 2019).

However, abundance-based analyses commonly assess taxa abundance at genus- or species-level, and the interaction of genetic diversity with BA metabolism within species has not yet been properly addressed. Since the functionality of a considerable proportion of microbial genes is still unknown (Heintz-Buschart and Wilmes, 2018), the homologue-based method for microbial BA gene analysis, which relies on the known references of BA biotransformation genes, has limited our understanding of the interactions of BAs with the “dark matter” of the gut microbiome. In addition, the accuracy of *in silico* modeling of BA biotransformations is affected by the possibility of undiscovered pathways in BA modification. Importantly, BAs themselves also influence gut microbiome composition through their antimicrobial activities and via indirect signaling pathways (Jia et al., 2017). However, the overall genetic shift of gut bacteria due to their exposure to the various BAs present in the human BA pool is currently unknown. This motivated us to explore the relationships between the gut microbiome and host BA metabolism at the level of microbial genetics.

Microbial structural variants (SVs) are highly variable segments of bacterial genomes that have been defined in recent years based on metagenomic sequencing data (Zeevi et al., 2019). Microbial SV regions potentially contain functional genes involved in host□microbe interactions and could thus provide information on sub-genome resolution of bacterial functionality. A variety of associations have been found between SVs and metabolite levels in human blood (Zeevi et al., 2019). Recently, a longitudinal study comparing subjects with irritable bowel syndrome to healthy individuals reported associations between BAs and microbial SVs for the first time (Mars et al., 2020). In this study, fecal levels of two unconjugated primary BA species, CDCA and CA, were found to correlate with variable genomic segments of *Blautia wexlerea*. This finding provided the initial clue that previously unknown bacterial genes are involved in the modification of primary BAs or indirectly associate with host BA metabolism (Mars et al., 2020). However, in view of the limited sample size and number of individual BA species analyzed in this study and the unknown reproducibility of the associations between SVs and BAs across different cohorts, systematic analysis in large-scale, population-based cohorts is required. Moreover, although the BA-associated SVs were interpreted as potential BA-metabolizing genomic segments (Mars et al., 2020), the existence of a causal relationship between BAs and microbial variants remains to be established because BAs can also act as regulators of the gut microbiome.

We therefore aimed to systematically evaluate the relationships between several parameters of human BA metabolism and the genetic architecture of the gut microbiome based on SVs. This study involved 1,437 individuals from two independent Dutch cohorts: the population-based Lifelines-DEEP cohort (LLD, *N* = 1,135) (Tigchelaar et al., 2015) and the 300-Obesity cohort (300-OB, *N* = 302) (Horst et al., 2019). In both cohorts, we profiled fasting plasma levels of 15 different BA species and 7α-hydroxy-4-cholesten-3-one (C4), which reflects the hepatic synthesis rate of BAs. We also calculated the relative proportions of individual BAs as well as different BA concentration ratios that represent metabolic pathways and enzymatic reactions. In all, we obtained 39 BA-related parameters. Simultaneously, the metagenomics sequencing data was subjected to characterization of microbial SVs to generate variable SV (vSV) and deletion SV (dSV) profiles that represent the standardized coverage and presence/ absence status of genomic segments, respectively. We then performed a systematic microbial genetic association analysis of BAs, not only with individual SVs, but also with discrete strains and the continuous genetic structures defined by the SV profiles. We further integrated several lifestyle factors, including diet, drug usage and smoking, and constructed tripartite networks of *in silico-*inferred causal relationships that included exposures, microbial genetics and host plasma BA composition. This identified potential novel microbial genetic regulators that mediate the effect of lifestyle on BA metabolism, which supports the potential of targeting the gut microbiome to alter human BA metabolism.

## Results

### High variability of plasma BA composition between individuals and cohorts

We included 1,437 individuals from two independent Dutch cohorts in this study: 1,135 individuals from the population-based LLD cohort and 302 individuals from the obese elderly-targeted 300-OB cohort (**Figure 1A-C**; **Table S1**). We assessed the concentrations and proportions of 15 BA species (6 primary and 9 secondary BAs) in fasting plasma: cholic acid (CA), chenodeoxycholic acid (CDCA), lithocholic acid (LCA), deoxycholic acid (DCA), ursodeoxycholic acid (UDCA) and their glycine- or taurine-conjugated forms (**Table S2**). We also computed 8 ratios that reflect hepatic and bacterial enzymatic activities (**Table S2**; **STAR**⍰**Methods**) and quantified the plasma level of C4, a biomarker of hepatic BA biosynthesis (Chiang, 2017). In total, we obtained 39 plasma BA parameters in this study.

**Figure 1.**
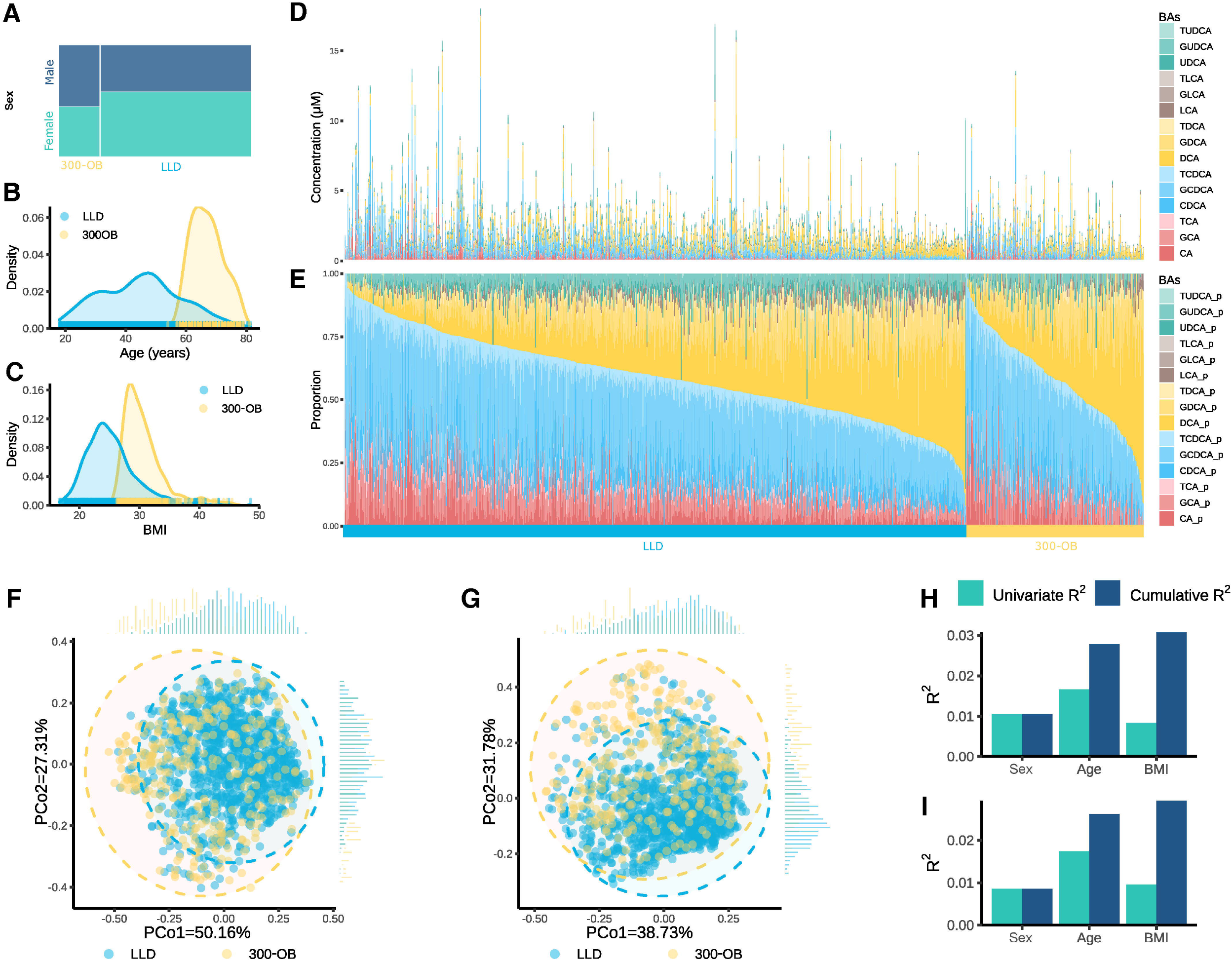
High variability in human fasting plasma bile acid concentration and composition. **A**. Sex proportions of LLD and 300-OB. **B**. Age distribution in LLD and 300-OB. **C**. BMI distribution in LLD and 300-OB. **D**. Concentrations of 15 bile acids (BAs) in fasting plasma across all samples of LLD and 300-OB. **E**. Proportions of 15 BAs in plasma across all samples of LLD and 300-OB. Samples were sorted by the proportion of the primary BAs (cholic and chenodeoxycholic acid and their conjugated forms) within each cohort. The order of samples is identical in (**D**) and (**E**). **F-E**, Principal coordinates analysis (PCoA) plot of the differences between all samples based on BA concentration profile (**F**) and BA proportion profile (**E**). **H-I**, Explained variance proportions (R^2^) of BA concentration (**H**) and proportion (**I**) profiles by sex, age and BMI. Blue bars indicate the cumulative explained BA variance proportion in multivariate models. Green bars indicate individually explained BA variance proportion by each factor in univariate models.

Both the concentrations and proportions of the 15 BA species showed considerable inter-individual variation in both cohorts (**Figure 1D** and **1E**). For instance, the total plasma BA concentration ranged from 0.084 to 18.008 μM (**Figure 1D**; **Table S3**) and the secondary/primary BA ratio ranged from 0 to 10.13 across all samples (**Figure 1E**; **Table S3**). Principal coordinate analysis (PCoA) also showed significant differences in plasma BA composition between LLD and 300-OB (Permutational multivariate analysis of variance (PERMANOVA), P = 0.001 in BA concentration profile, P = 0.001 in BA proportion profile; **Figure 1F** and **1G**). We also observed that 34 of the 39 BA parameters showed significant differences between LLD and 300-OB (Wilcoxon rank-sum test, FDR < 0.05; **Figure S1A**; **Table S4**). In view of the distinctly different characteristics of participants between LLD and 300-OB (**Figure 1A-1C**), the difference in BA composition between the two cohorts could be caused by phenotypic differences. We therefore estimated the explanatory power of basic phenotypes such as age, sex and body mass index (BMI) on the variation in BA composition. These factors collectively explained only 3.07% and 2.94% of the variance in plasma BA concentration and BA proportion profiles, respectively (**Figure 1H** and **1I**). Here, age had the largest effect, explaining 1.66% and 1.74% of the variance in plasma BA concentration and proportion profiles, respectively (PERMANOVA, p < 0.05; **Figure 1H** and **1I**). This suggests that a large proportion of BA variation remains unexplained and may be attributed to other factors such as lifestyle factors, host genetic background and gut microbial factors.

### SV profiling unravels microbial genetic differences between the general population-based and the obesity-based cohorts

Using the metagenomics sequencing data from both cohorts, we detected 8,282 SVs, including 2,616 vSVs and 5,666 dSVs in 55 species that were present in at least 75 samples in the two cohorts with sufficient coverage across the reference genomes (**STAR**⍰**Methods**) with 32–374 SVs per species (**Figure 2A** and **2B**; **Table S4**). These 55 species together accounted, on average, for 66.60% of the total microbial species composition, ranging from 27.02%–90.25% (**Figure S2A**). The average sample size of all 55 species with SVs was 432 (**Figure S2B**; **Table S5**). The most prevalent species with SV calling was *B. wexlerae*, which could be detected in 1,350 samples (1,071 from LLD and 279 from 300-OB), followed by *E. rectale* (*N* = 1,160), *E. hallii* (*N* = 1,138) and *Ruminococcus* sp. (*N* = 1,095).

**Figure 2.**
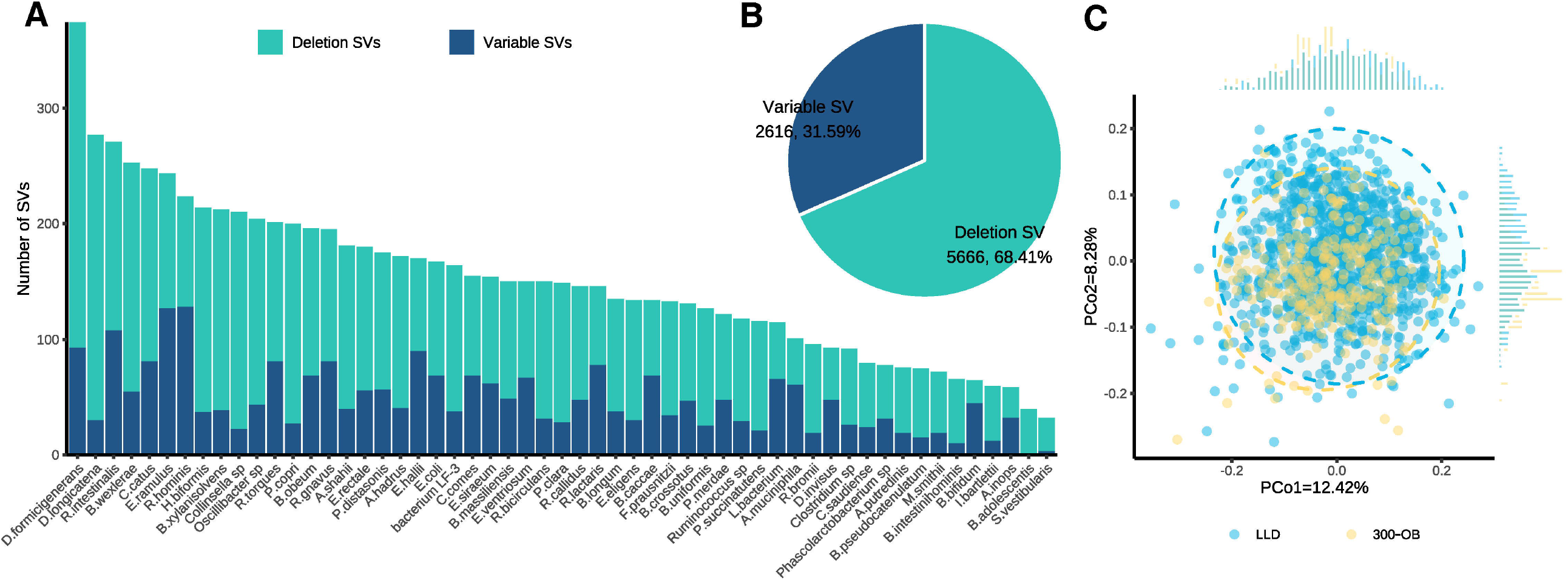
Overview of structural variation profile in LLD and 300-OB. **A**. Number of structural variants (SV) of each species. **B**. Total SV numbers. **C**. Population structure of SV-based genetic makeup.

We further assessed the Canberra distance of bacterial SV profiles between all samples (**Figure 2C**). Principal components (PCo) 1 and 2 together explained 20.70% of the total SV-based genetic variance (**Figure 2C**), which showed significant differences between LLD and 300-OB (Wilcoxon rank-sum test, P = 9.83×10^−4^ for PCo1 and P = 2.62×10^−11^ for PCo2), indicating the divergence of microbial genetics between the general population-based cohort and the obesity cohort. Interestingly, age, gender, BMI and read counts collectively only explained 1.79% of the variance of the metagenome-wide SV profile (**Figure S2C**), while the top factor, total read counts, explained only 0.7% of the variation.

### Species-level genetic makeup correlates with human BA metabolism independent of bacterial species abundance

We first investigated the taxonomic abundance and microbial genetic associations with fasting plasma BA parameters separately at the species level (**Figure S3A** and **S3B**). In total, we identified 226 significant associations between the relative abundance of 34 bacterial species and 36 BA parameters (Linear regression, FDR_Meta_<0.05; **Figure S3B**; **Table S6**). Several BA parameter-associated species had been reported earlier in the 300-OB cohort, e.g. the negative association of the relative abundance of the butyrate-producing species *F. prausnitzii* with 20 BA parameters, including a negative association with secondary/primary BA ratio (Linear regression, Beta_Meta_=-0.18, FDR_Meta_=3.97×10^−6^; **Table S6**). The relative abundance of another butyrate-producing species, *E. hallii*, correlated with C4 concentration (Linear regression, Beta_Meta_=0.10, FDR_Meta_=4.72×10^−3^; **Table S6**), consistent with findings from a mouse study showing that *E. hallii* is able to modify BA metabolism (Udayappan et al., 2016). The most significant abundance association was found between *R. gnavus* and UDCA proportion in plasma (Linear regression, Beta_Meta_=0.34, FDR_Meta_=7.77×10^−28^; **Table S6**). Altogether, these results confirm that microbiome composition is closely associated with human BA metabolism.

As bacterial genomes are highly variable, the microbial genetic content of each species varies across different individuals (Rossum et al., 2020; Tierney et al., 2019), which may also be relevant to human BA metabolism. Therefore, we first calculated the SV-based genetic distance per species (**Figure S4**) and associated these with the 39 plasma BA parameters using PERMANOVA, after correcting for age, sex, BMI, read counts and species abundance if applicable (**STAR**⍰**Methods**). In total, we identified 260 significant associations between genetic distances of 39 bacterial species and 36 BA parameters (PERMANOVA, FDR_Meta_ < 0.05; **Figure S3A**; **Table S6**), which indicates that SV-represented microbial genetic associations with BA parameters are largely independent of the relative abundances of the species. Interestingly, some species were found to be more likely associated with BA parameters at the genetic level (e.g. *C. comes, E. rectale* and *R. intestinalis*), whereas other species tended to be associated with BA parameters at the relative abundance level (e.g. *A. muciniphila, B. bifidum, B. crossotus* and *I. bartlettii*) (**Figure 3A**). Out of 260 BA associations with species-specific genetic makeup, only 50 were also detected at the species abundance level (**Figure S3C**; **Figure 3A**), which highlights that microbial genetic variation represents a new layer of information about bacterial functionality.

**Figure 3.**
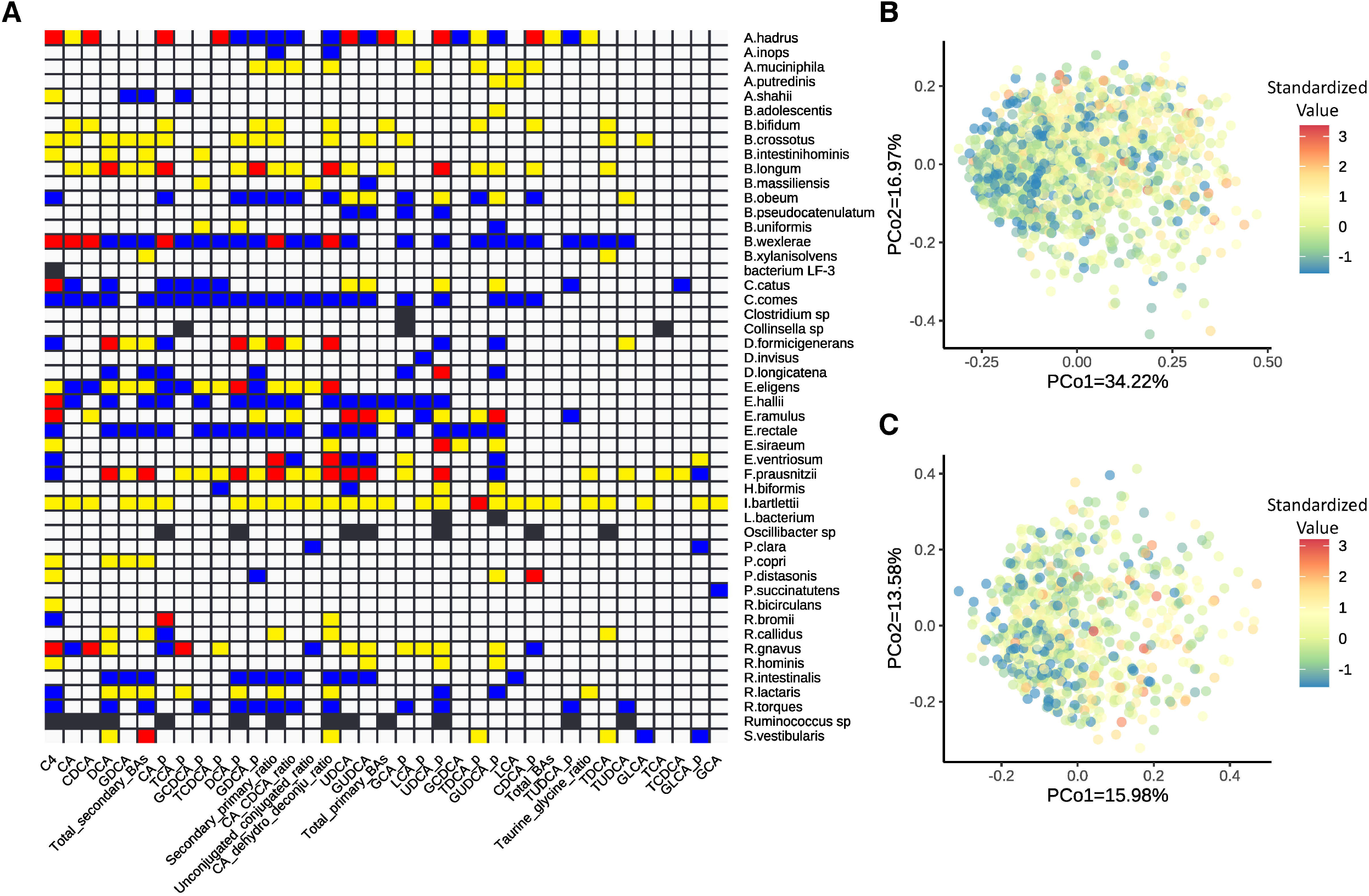
Species-level associations of gut microbiome with human bile acid parameters. **A**. Heatmap of species-level associations with BA parameters. Blue indicates purely genetics-based associations. Yellow indicates purely relative abundanceLbased associations. Red indicates associations based on both genetics and relative abundance. Black indicates genetics-based associations where relative abundance is not available for the corresponding species. White indicates no association. **B**. Genetic association of *B. wexlerae* with CA proportion in plasma, the color scale from red to blue represents the increase in standardized value of CA proportion. **C**. Genetic association of *F. prausnitzii* with GUDCA proportion in plasma the color scale from red to blue represents the increase in standardized value of GUDCA proportion.

The species with the highest number of genetic associations was *B. wexlerae*. The inter-individual genetic differences of *B. wexlerae* were significantly associated with 27 BA parameters (PERMANOVA, FDR_Meta_<0.05; **Table S6**), whereas only 6 BA parameters correlated with the relative abundance of *B. wexlerae* (Linear regression, FDR_Meta_<0.05; **Table S6**). The strongest genetic association of *B. wexlerae* was with plasma CA proportion (P_Meta_ = 8.70×10^−6^; **Figure 3B**; **Table S6**), indicating that individuals with very similar *B. wexlerae* genome content tend to have similar CA-contributions to their plasma BA content. Another species, *F. prausnitzii*, contributes to 12-dehydro-CA production, and the depletion of *F. prausnitzii* was inferred to lower the unconjugated CA and CDCA levels in feces of IBD patients (Heinken et al., 2019). In addition to associations at species-abundance level, genetic differences in *F. prausnitzii* were also associated with 23 BA parameters (**Table S6**). For instance, genetic differences in *F. prausnitzii* correlated with the proportion of GUDCA in plasma (PERMANOVA, FDR_Meta_<0.05; **Figure 3C**; **Table S6**), even though the association was not significant at species-abundance level. Altogether, we observed that species-specific genetic makeup is highly variable and correlates with BA composition independent of the relative abundances of the species in the microbial community.

### Discrete subspecies correlate with human BA metabolism

Based on the genetic differences between species, we stratified the population genetic structure for each species using the partitioning around medoid□based method (**STAR**⍰**Methods**; **Figure S4**) and detected two or more subspecies for 29 of the 55 species (**Figures S5** and **S6**; **Table S7**). Some subspecies have been previously reported based on different methods. For instance, we identified two *E. rectale* subspecies and four *A. muciniphila* subspecies in our cohorts, while Costea *et al*. observed three *E. rectale* and two *A. muciniphila* subspecies based on a single-nucleotide variant (SNV)-typing profile in 2,144 samples (Costea et al., 2017). All the subspecies we identified could be detected in both LLD and 300-OB samples, but the subspecies proportions of *P. copri, S. vestibularis* and *P. merdae* showed different enrichments in LLD and 300-OB (chi-square test, FDR < 0.05). Of these, *P. copri* subspecies 1, *S. vestibularis* subspecies 1 and *P. merdae* subspecies 2 were enriched in LLD, whereas *P. copri* subspecies 2, *S. vestibularis* subspecies 3 and *P. merdae* subspecies 1 were enriched in 300-OB (Chi-square test, FDR < 0.05; **Figure S7**; **Table S8**). Consistent with the previous results reported for SNV-based subspecies of *E. rectale* (Costea et al., 2017), the SV-based subspecies of *E. rectale* we identified was also associated with host BMI (Wilcoxon test, P = 0.0045).

The presence of different subspecies may differentially affect BA metabolism. We therefore conducted an association analysis between the SV-based subspecies and plasma BA parameters and found 41 significant associations (Permutational Kruskal-Wallis rank-sum test, FDR < 0.05; **Figure 4**; **Table S9**). *E. rectale* showed the highest number of associations with BAs (10 associations), followed by *R. gnavus* (8 associations). The most significant association was between *E. rectale* and C4 concentration (Permutational Kruskal-Wallis rank-sum test, FDR = 2.26×10^−5^). We compared the SV profiles of two subspecies of *E. rectale* and found that 55 of the 56 vSVs and 72 of the 124 dSVs were significantly enriched in subspecies 1 or 2 (Wilcoxon test for vSV and Chi-square test for dSV, FDR < 0.05). Although no enrichment was observed for BA biotransformation genes between *E. rectale* subspecies, their wide association with BA parameters may reflect a physiological impact of bacterial subspecies diversity on BA metabolism, or *vice versa*. The antimicrobial activity of BAs exerts survival pressure on microbes (Langdon et al., 2016; Tian et al., 2020), which may drive some or all of the changes in bacterial genomes.

**Figure 4.**
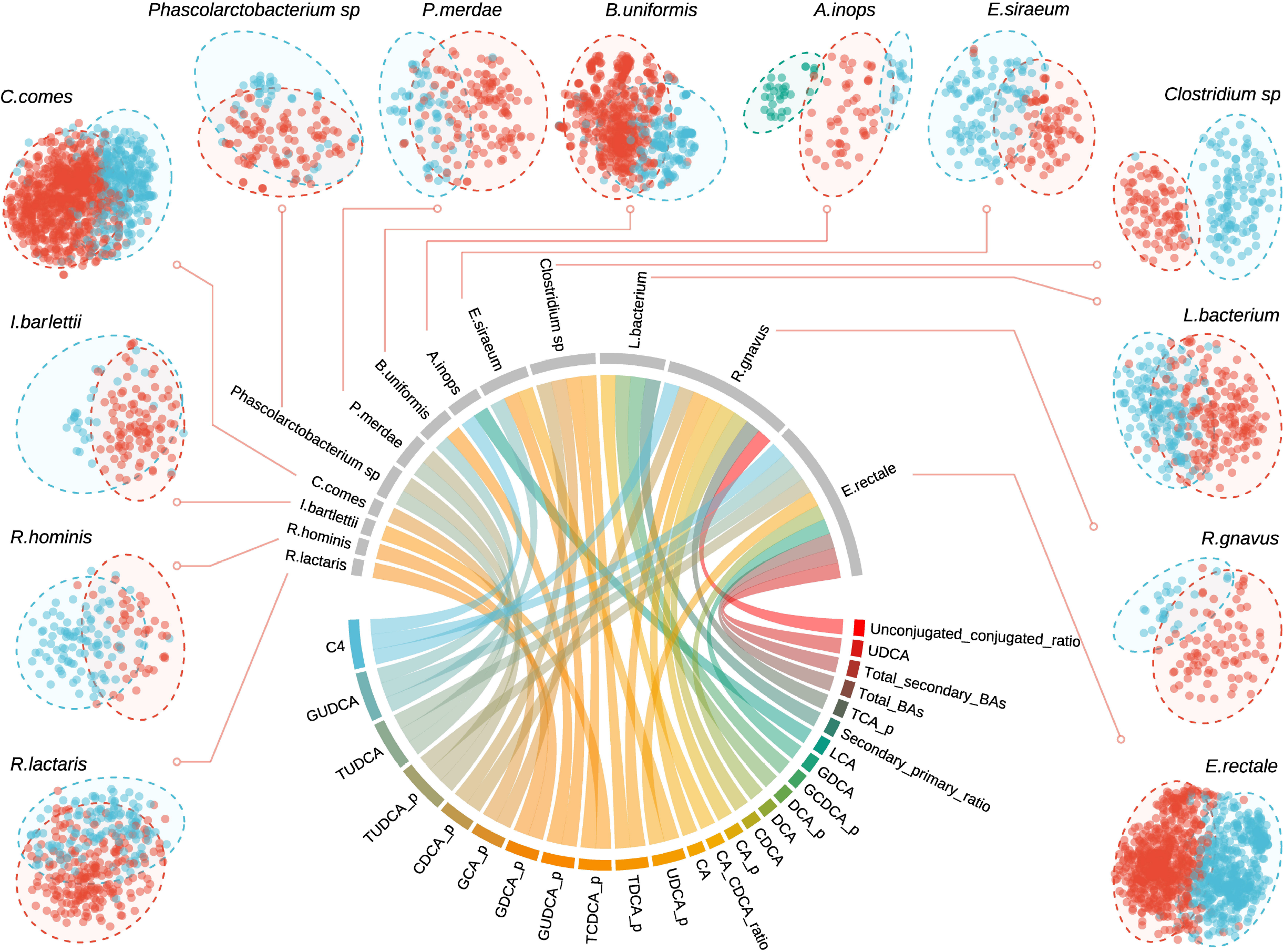
Bile acid parameters correlate with structural variant□based subspecies. The subspecies of 13 species are shown by the t-SNE plots, with distinct subspecies shown by different colors. The circos correlation plot shows their associations with BA. Each line indicates an association between subspecies of a species and plasma levels of a BA parameter.

### Metagenome-wide SV-based association identifies BA-associated microbial genomic segments

To identify SVs that potentially harbor genes involved in human BA metabolism, we performed a metagenome-wide microbial SV-based association study on the BA parameters. Considering the significant differences in plasma BA composition and microbial genetic makeup between LLD and 300-OB, we associated the 8,282 SVs with the 39 plasma BA parameters using linear models for LLD and 300-OB respectively, followed by meta- and heterogeneity analysis (random effect model). In addition to age, sex, BMI and total read counts, we also included the corresponding species abundances as a covariate because we observed that the abundance levels of 34 of the 55 species were associated with at least one BA parameter (**Table S6**). In total, we identified 792 significant and replicable associations in our meta-analysis using a random effect model (FDR_meta_<0.05), including 725 associations with vSVs (**Figure S8A**; **Table S10**) and 67 associations with dSVs (**Figure S8B**; **Table S11**). The effect sizes and directions of all 792 associations were highly consistent between cohorts (P_hetero_<0.05; **Figure S8C** and **S8D**). These results indicate that the SV associations we identified were robust and replicable between the two cohorts despite the large differences in their profiles of gut microbial genetic makeup and plasma BA composition.

The 792 replicable SV-BA associations linked 300 SVs of 33 species with 32 plasma BA parameters **(Figure 5A**), indicating that BA-related SVs are highly prevalent across gut bacterial species. 183 (23.11%) of the 792 replicable associations were for *B. wexlerae* (**Figure 5B**). In the genome of *B. wexlerae*, 52 SVs were associated with 21 BA parameters (FDR_Meta_<0.05; **Figure 5B**), with the most significant association being between the variable SV region 1715-1716 kbp and the CA dehydroxylation/deconjugation ratio (Beta = -0.29, P_meta_ = 2.18×10^−23^; **Table S10**). Since the reference genome of *B. wexlerae* was not well annotated in the database provided by SGVFinder, we further annotated its genome with PATRIC (**STAR**⍰**Methods**) and identified three genes that encode choloylglycine hydrolase (or bacterial BSH; EC number: EC 3.5.1.24), which catalyzes the deconjugation of glycine- and taurine-conjugated BAs. One of the annotated BSH genes is located in the region of 2,938,104–2,938,946 bp, which is close to three BA-associated SVs of *B. wexlerae* that span four genomic segments (2,079–2,081 kbp, 2,081–2,082 kbp, 2,083–2,084, and 2,086–2,090 kbp) (**Figure 5B** and **5C**). These three SVs also significantly correlated with 10 BA parameters (FDR_Meta_ < 0.05; **Figure 5B**), and the most significant association was with DCA concentration (Beta_Meta_ = -0.23, P_Meta_ = 1.67×10^−16^; **Figure 5D**; **Table S10**). This is in line with recent findings that fecal CDCA and CA were associated with several SVs of *B. wexlerae* (Mars et al., 2020). Our study further confirms that *B. wexlerae* is a novel bacterium involved in BA transformation that possesses BA metabolism□related genes.

**Figure 5.**
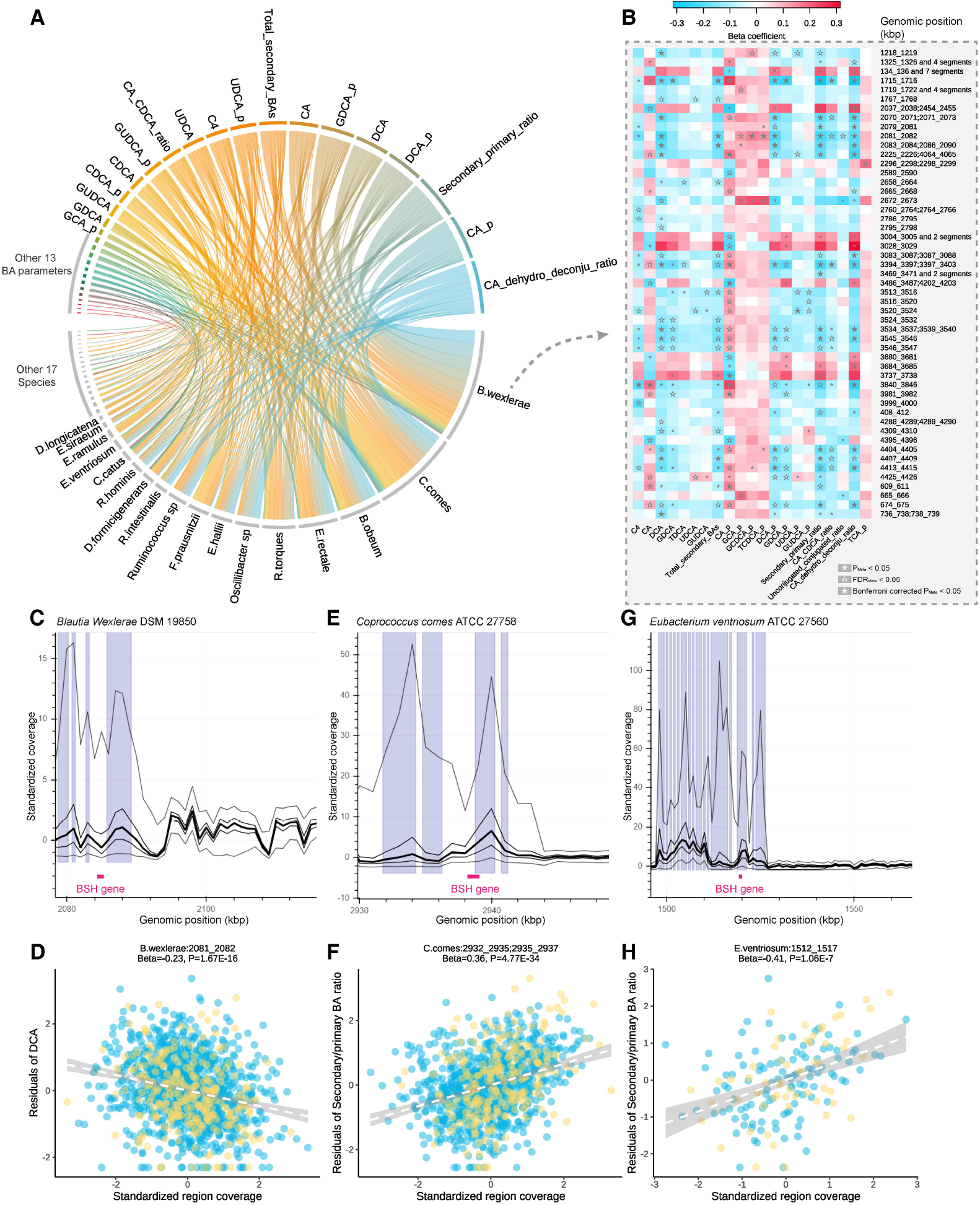
Associations between bile acid parameters and structural variants. **A**. Replicable significant associations between BA parameters and SVs (FDR_Meta_<0.05). **B**. Heatmap of associations between BA parameters and SVs of *Blautia wexlerae*. **C-H**. Examples of SV regions close to known BA biotransformation genes (**C, E**, and **G**) and associations with BA parameters (**D, F**, and **H**). Blue and yellow circles represent LLD and 300-OB samples, respectively.

Besides *B. wexlerae*, we also identified BA biotransformation genes near the associated SV regions in several other species. For instance, a 5-kbp vSV (2,932□2,935 and 2,935□2,937 kbp) of *Coprococcus comes* harbors a BSH gene (2,938,104–2,938,946 bp) that was significantly associated with 12 BA parameters (**Figure 5E**; **Table S10**), of which the strongest association was with the secondary/primary BA ratio (Beta_Meta_ = 0.36, P_Meta_ = 4.77E×10^−34^; **Figure 5F**). This appeared to be the most significant association among all SV-BA associations. In addition to being found in the most prevalent species, BA biotransformation genes were also found in some low prevalence species. For instance, near a BSH gene (genomic position: 1,519,343□1,520,332 bp) in the genome of *E ventriosum*, three variable SVs were significantly associated with 10 BA parameters (FDR_Meta_ < 0.05) (**Figure 5G**; **Table S10**), of which the most significant association was between the vSV region 1,512□1,517 kbp and the secondary/primary BA ratio (Beta_Meta_ = -0.41; P_Meta_ = 1.06×10^−7^; **Figure 5H**).

We also identified 118 BA□SV associations with significant heterogeneity between our general population- and obesity-based cohorts (P_hetero_ < 0.05, FDR_LLD_ < 0.05 and/or FDR_300-OB_ < 0.05; **Table S12** and **S13**). The most significant heterogeneity was observed for the association between a 3-kbp vSV of *Escherichia coli* (1,062□1,065 kbp) and TCDCA proportion (Beta_LLD_ = -0.16, FDR_LLD_ = 0.76, Beta_300-OB_ = 0.68, FDR_300-OB_ = 0.032, P_hetero_ = 3.24×10^−6^; **Table S12**). This variable genomic region contains two genes, *Salmochelin siderophore protein IroE* and *Enterochelin esterase*, that play a role in maintaining iron homeostasis of *E. coli*. A 1-kbp vSV of *C. comes* (966 □967 kbp) harboring a BSH gene was associated with three BA parameters (CA/CDCA ratio, secondary/primary BA ratio and DCA proportion) with significant heterogeneity between LLD and 300-OB (P_hetero_ < 0.05; **Table S12**), and the effect sizes of the BA associations were higher in 300-OB than in LLD.

### Bi-directional causality between bacterial SVs and host BAs

The genetic makeup of gut bacteria can be affected by host-specific features and environmental factors, thus a symbiotic genomic variant □based GWAS alone will not be sufficient to provide causal interpretation of the relationships between correlated microbial variants and host phenotypes. Although we did identify several bacterial genes known to be involved in BA biotransformation that were located in BA-associated SV regions, the causality behind most of the BA □SV associations we identified remains unknown. The lifestyle exposure factors collected in the LLD cohort enabled us to infer *in silico* causal relationships between correlated SVs and BAs and identify lifestyle factors that may impact the interactions between gut microbial genetics and host BA metabolism. We integrated 127 lifestyle factors (78 dietary factors, 44 drug usage factors and 5 smoking-related factors; **Table S14**) with SV and BA data. Here, we first identified lifestyle □SV □BA groups in which all the variables correlated with each other and then conducted bidirectional mediation analysis. In the first causal direction, we hypothesized that SVs act as regulators that mediate the effects of lifestyle factors on the composition of the BA pool and thus treated SVs as mediators and BA parameters as outcomes. In the second causal direction, we assessed whether BAs can mediate the effects of lifestyle factors on bacterial SVs (**Figure 6A**). In total, we identified 502 groups of inferred *in silico* causal relationships, including 38 unidirectional causal relationships in direction 1, 216 unidirectional causal relationships in direction 2 and 248 bidirectional causal relationships (**Figure 6B**, FDR_mediation_ < 0.05).

**Figure 6.**
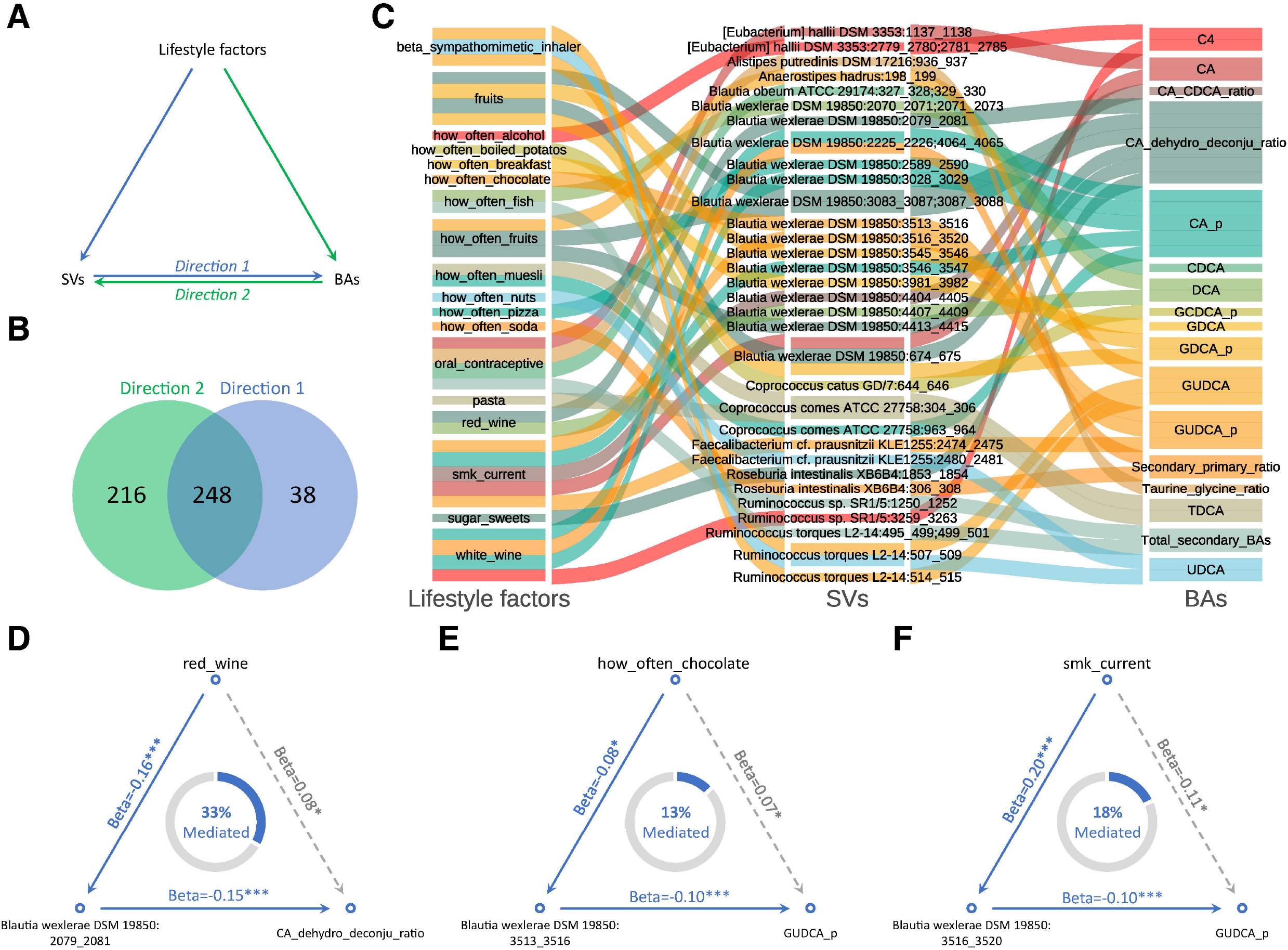
Causal relationship inference using bi-directional mediation analysis. **A**. Framework of bi-directional mediation analysis between lifestyle factors, SVs and BAs. **B**. Number of inferred causal relationships for direction 1 (from SV to BA), direction 2 (from BA to SV), and both. **C**. Sankey diagram showing the inferred causal relationship network of direction 1. **D-F**. Examples of causal relationships between lifestyle factors, SVs and BAs inferred by bi-directional mediation analysis. The beta coefficient and significance are labeled at each edge and the proportions of indirect effect (mediation effect) are labeled at the center of the ring charts.

The tripartite causal network in direction 1 was composed of 18 lifestyle factors, 32 SVs as mediators and 17 plasma BA parameters as outcomes (FDR_mediation_ < 0.05, **Figure 6C, Table S15**). Notably, 15 of the 32 SVs were from *B. wexlerae*, including the SVs with known BA biotransformation genes. For instance, a 2-kbp vSV (2079□2081 kbp) close to a BSH gene in *B. wexlerae* regulated the effect of drinking red wine on the CA dehydroxylation/deconjugation ratio (FDR_mediation_ < 0.05; Mediated proportion = 33%; **Figure 6D**). Another SV of *B. wexlerae* in 3513□3516 kbp mediated both the positive effect of the frequency chocolate consumption on GUDCA proportion in plasma (FDR_mediation_ < 0.05; Mediated proportion = 13%; **Figure 6E**) and the inverse effect of smoking habit on this parameter (FDR_mediation_ < 0.05; Mediated proportion = 18%; **Figure 6F**). These findings indicate that gut bacterial genes involved in BA metabolism can be regulated by lifestyle factors and thereby affect the composition of the host’s BA pool. The inferred regulatory SVs that causally affect the composition of host BA pool may contain novel genes involved in BA biosynthesis and biotransformation and can potentially be used as targets to regulate BA metabolism.

We found 216 *in silico* causal relationships in which 22 BA parameters mediated the effects of 43 lifestyle factors on 80 bacterial SVs belonging to 12 bacterial species (FDR_mediation_ < 0.05; **Table S15**). Among the 80 regulated SVs, 29 were from *B. wexlerae*, followed by 11 from *R. torques*. In mice, the growth of *Ruminococcus* species can be inhibited by DCA (Tian et al., 2020), and we found 8 SVs of *Ruminococcus* species to be negatively regulated by both DCA concentration and proportion (FDR_mediation_ < 0.05), indicating that enrichment of the circulating BA pool with DCA may exert selective pressure on *Ruminococcus* species and cause a loss of their genomic content.

## Discussion

We characterized the gut microbial SV and plasma BA profiles of 1,437 Dutch individuals from two independent cohorts and systemically assessed the correlation between gut microbial genetics and host BA metabolism from species genetic makeup level down to single variant level. Species genetic makeup was found to correlate with BA parameters independent of the relative abundances of these species. We also identified subspecies of 29 bacterial species using SV-based clustering analysis, revealed the within-species genetic differentiation and diversity and associated the SV-based subspecies with plasma BA parameters. We further performed a metagenome-wide microbial SV association study on 39 BA parameters and identified 786 replicable associations between SVs and BA parameters and 118 heterogeneous associations using meta-analysis. Bi-directional mediation analysis inferred *in silico* regulatory relationships behind the correlations we identified. Our study thus provides a resource of bacterial genomic entities that potentially contain novel genes involved in human BA metabolism while also revealing genetic shifts in the bacterial genomes that are potentially due to the antimicrobial effects of specific BAs within the intestinal lumen. To the best of our knowledge, this is the largest study so far on microbial genetic determinants of plasma BA concentrations and composition in humans. In view of the growing awareness of the involvement of specific BAs in the onset and progression of human diseases (Chávez-Talavera et al., 2017; Dermadi et al., 2017; Gao et al., 2019), as well as the current development of pharmacological agents that target BA-signaling pathways for treatment of liver and metabolic diseases (Jia et al., 2017; Krautkramer et al., 2021; Pathak et al., 2018; Sun et al., 2018), this knowledge is of direct clinical relevance.

Our study demonstrates that SV-based metagenome-wide association is a powerful method to bring microbial associations closer to functionality and mechanistic understanding. Firstly, our study shows that the BA associations with microbial SVs were often stronger than those with species relative abundances and can even be independent of species relative abundances. This highlights the value of metagenomic SVs as a new source of information that describes the functionality of the human gut microbiome. Although previous studies identified some BA-metabolizing species (Krautkramer et al., 2021; Li et al., 2021), our metagenome-wide SV association study supplements the list of novel bacterial species that interact with BA metabolism, adding *Blautia wexlerae, Eubacterium rectale, Blautia obeum* and *Ruminococcus torques*, amongst others. The sub-genome scale analysis also pinpoints the location of genomic segments that associate with host BA pool, which means that association of microbial SV across the whole metagenome with host phenotypes helps to locate microbial genes or genetic elements involved in hostLmicrobe interaction. Our study underscores the contribution of gut microbial genetics to the individuality of host BA metabolism. The comprehensive association analysis approach we used provides a template for cohort-based microbial genetics studies, demonstrating a paradigm shift from “micro-ecology” to “micro-population genetics”.

Our study further highlights the complex, bi-directional effect between the gut microbiome and BA metabolism. We used lifestyle factors as exogenous predictors to infer *in silico* potential causal relationships between SVs and BAs using bidirectional mediation analysis and identified specific lifestyle factors involved in the interaction between bacterial genetics and BA metabolism. This highlights the potential of targeting the gut microbiota to regulate BA metabolism through lifestyle intervention. For instance, we found that an SV of *B. wexlerae* mediated the effect of red wine drinking on the CA dehydroxylation/deconjugation ratio, reflecting the conversion of glycine- or taurine-conjugated CA to unconjugated DCA within one cycle of the enterohepatic circulation. Red wine is rich in polyphenols, a group of molecules with anti-oxidative properties (Naumann et al., 2020; Queipo-Ortuño et al., 2012) that can increase fecal BA excretion by regulating gut microbiota (Chambers et al., 2019). We also observed that the frequency of chocolate consumption increases the GUDCA proportion in plasma through an SV of *B. wexlerae*. GUDCA is a hydrophilic BA that has been suggested to act as an antagonist of human FXR and to contribute to the beneficial effects of metformin in subjects with T2D (Sun et al., 2018). Furthermore, its parent molecule, UDCA, is widely used in treatment of cholestatic liver diseases and has been suggested as a potential drug for the treatment of T2D and other metabolic diseases (Pathak et al., 2018; Sun et al., 2018). Chocolate is rich in flavonoids, a subclass of polyphenols. Thus, it appears that polyphenols from chocolate increase the level of GUDCA by regulating gut bacterial genes. Previous studies reported that dietary polyphenols from plant-derived foods can affect the composition of fecal BAs in humans by regulating gut microbiota (Chambers et al., 2019; Ozdal et al., 2016; Queipo-Ortuño et al., 2012; Sembries et al., 2006). Our *in silico* causal inference analysis revealed that the bacterial SV serves as a mediator that regulates the effects of dietary polyphenols on BA metabolism. Conversely, our study also provided evidence that BAs, likely via their anti-bacterial activities as “intestinal soaps”, not only affect the growth of intestinal microbes but also pose selective pressure on bacterial genetics.

## Limitations of the study

We acknowledge several limitations of our current study. We investigated the association between plasma BA parameters and variable genomic segments of gut bacteria in two independent cohorts, identified substantial consistent associations in both these general population and obese individuals, and demonstrated the reliability of BA associations with microbial SVs. However, all the samples included in this study were collected from the residents of the Netherlands. Considering the potential heterogeneity of host □microbiome interaction across populations with different genetic and environmental backgrounds, the associations between plasma BA parameters and microbial SVs need to be replicated in other populations with different background. As this is a cross-sectional study, we inferred the regulatory relationships between BA parameters and microbial SVs using mediation analysis, but whether the shifts of microbial genetic elements causally correlate with host BA metabolism still requires further confirmation in a longitudinal study design and through experimental validation. Additionally, plasma BA parameters cannot fully represent the flux of the BA pool in enterohepatic circulation and are only modestly correlated with the fecal BA pool (Chen et al., 2020), further study of the association between microbial genetic variation and BA metabolism in their actual niche □ the enterohepatic circulation □ is thus needed. Despite these limitations, our study represents a step towards successful microbiome-targeted interventions to improve host metabolism, in particular through modulation of BA metabolism, which is a major target for the treatment of NAFLD and its metabolic co-morbidities.

## Supplemental information

Supplemental materials are available.

## Acknowledgements

We thank all the volunteers in the LifeLines-DEEP cohort and 300-Obesity cohort of the Human Functional Genomics Project (HFGP) for their participation and the project staff for their help and management. This study was supported by an IN-CONTROL CVON grant (CVON2012-03, CVON2018-27) to N.P.R., M.G.N., A.Z., F.K. and J.F. J.F. is supported by an ERC Consolidator grant (101001678) and the Netherlands Organ-on-Chip Initiative, an NWO Gravitation project (024.003.001) funded by the Ministry of Education, Culture and Science of the government of the Netherlands. F.K. is supported by the Noaber Foundation, Lunteren, the Netherlands. A.Z. is supported by ERC Starting Grant 715772, NWO-VIDI grant 016.178.056 and NWO Gravitation grant Exposome-NL (024.004.017). D.W. is supported by China Scholarship Council (CSC201904910478). L.C. holds a joint fellowship from the University Medical Centre Groningen and China Scholarship Council (CSC201708320268) and a Foundation de Cock-Hadders grant (20:20-13). M.D. holds a MD-PhD fellowship from the University Medical Center Groningen. We also thank the Genomics Coordination Center for providing data infrastructure and access to high performance computing clusters and Kate Mclntyre for critical reading and editing.

## Author contributions

J.F., F.K. and A.Z. conceptualized and managed the study. D.W., M.D., L.C., I.C.L.v.d.M., M.K., N.P.R. and J.H.W.R. contributed to sample collection and data generation. D.W. analyzed the data. D.W., J.F. and F.K. drafted the manuscript. D.W., M.D., L.C., S.A.S., I.C.L.v.d.M., H.A., M.K., V.W.B., N.P.R., J.H.W.R., M.G.N., A.Z., F.J. and F.K. reviewed and edited the manuscript.

## Additional information

### Lead contact

Further information and requests for resources, software, reagents and data sharing should be directed to the Lead Contact, Jingyuan Fu.

### Data and Code Availability

Raw metagenomic sequencing data of LifeLines-DEEP and 300-Obesity are publicly available at European Genome□Phenome Archive via accession numbers EGAS00001001704 and EGAS00001003508, respectively. The code used for the statistical analysis is available via https://github.com/GRONINGEN-MICROBIOME-CENTRE/Groningen-Microbiome/tree/master/Projects/SV_BA.

## STAR⍰Methods

## KEY RESOURCES TABLE

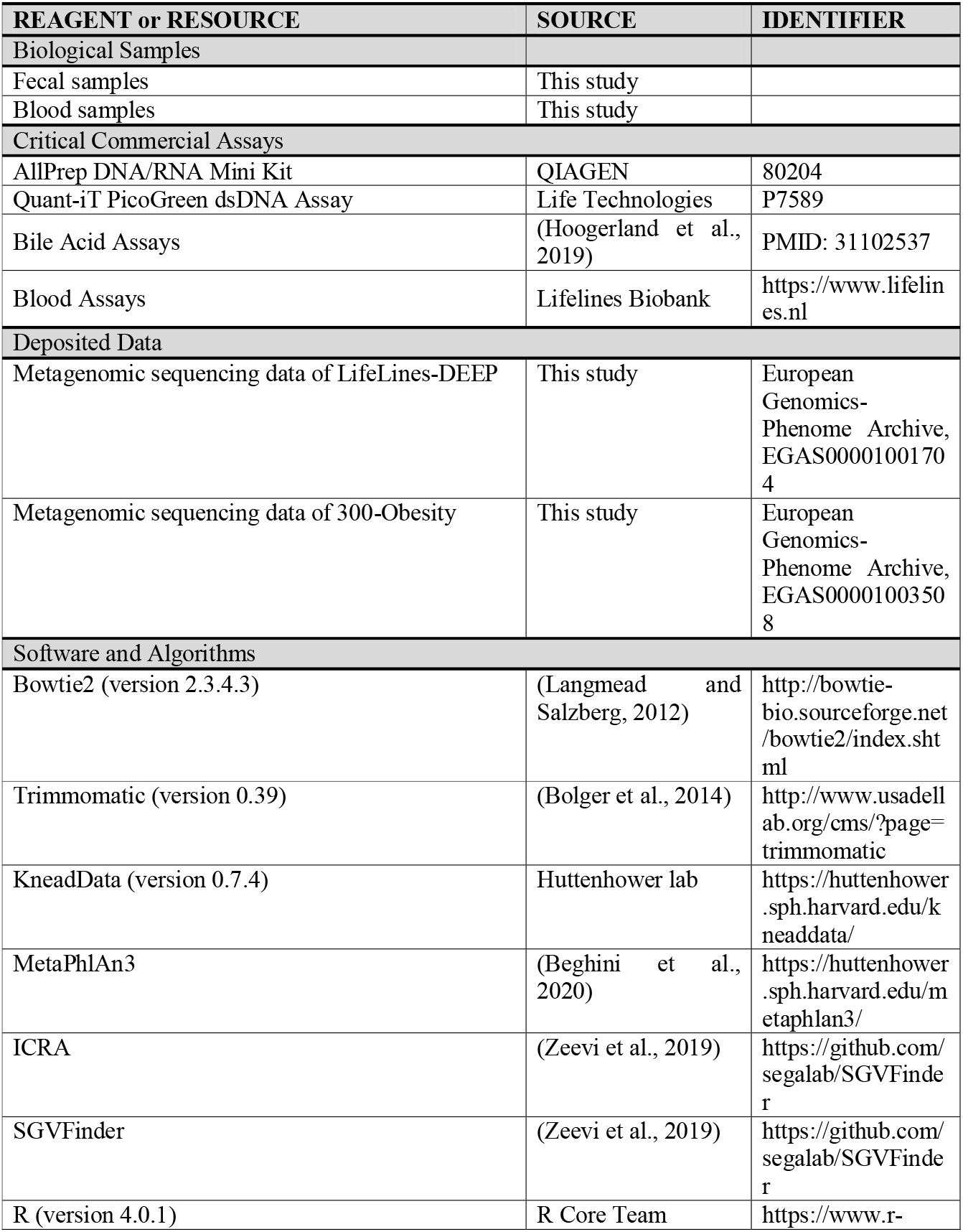
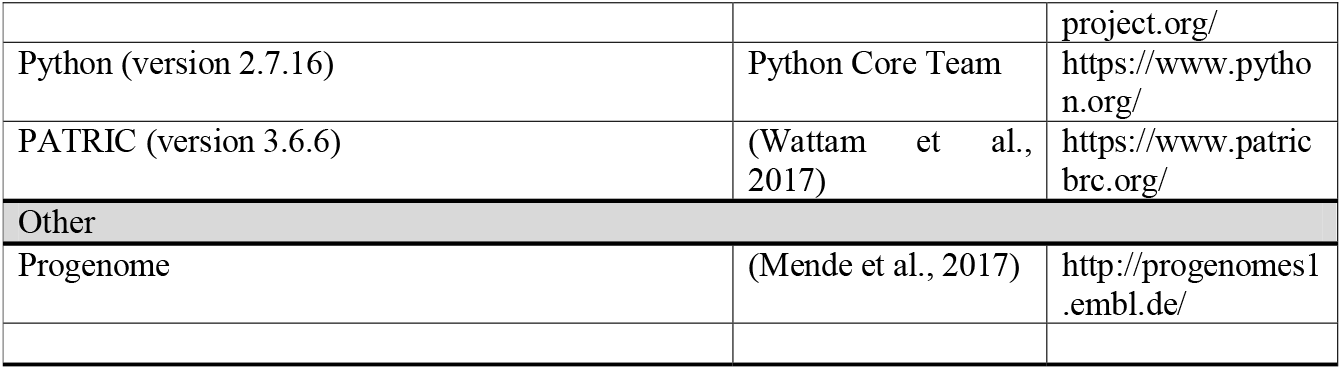

## RESOURCE AVAILABILITY

### Materials Availability

This study did not generate new unique reagents.

### Data and Code Availability

Raw metagenomic sequencing data of LifeLines-DEEP and 300-Obesity are publicly available from the European Genome-Phenome Archive via accession number EGAS00001001704 and EGAS00001003508, respectively. The code used for statistical analysis is available via https://github.com/GRONINGEN-MICROBIOME-CENTRE/Groningen-Microbiome/tree/master/Projects/SV_BA.

## EXPERIMENTAL MODEL AND SUBJECT DETAILS

### LifeLines-DEEP cohort

LifeLines-DEEP (LLD) is a sub-cohort of LifeLines, a large population-based prospective cohort that enrolled 167,729 participants from the north of Netherlands, established to explore the risk factors of complex diseases. In LLD, 1,539 individuals were included and multi-layers of omics data were collected. In the current study, high-quality metagenomic sequencing data, 78 dietary factors, 5 smoking factors and 44 drug usage factors were available for 1,135 individuals (474 males and 661 females). The average age of LLD participants was 45.04 years old (18□81, SE = 0.40) and the average BMI was 25.26 (16.67–48.56, SE = 0.12).

### 300-Obesity cohort

The 300-Obesity (300-OB) cohort was established by Radboud University Medical Center, Nijmegen, the Netherlands (Horst et al., 2019). In total, 302 individuals (167 males and 135 females) aged 55□81 years with a high body mass index (BMI) > 27 were enrolled in 300-OB. The average age of 300-OB participants was 67.1 years old (54□81, SE = 0.31) and the average BMI was 30.7 (26.3–45.5, SE = 0.20). All participants were included between 2014 and 2016.

### Ethical approval

The LifeLines-DEEP study has been approved by the Institutional ethics Review Board (IRB) of the University Medical Center Groningen (ref. M12.113965), the Netherlands. The 300-Obesity study has been approved by the IRB CMO Regio Arnhem-Nijmegen (nr. 46846.091.13).

## METHOD DETAILS

### Bile acid quantification

Levels of 15 BAs and C4 concentrations in fasting plasma were quantified by liquid chromatography–mass spectrometry (LC-MS) procedures, as previously described (Eggink et al., 2017; Hoogerland et al., 2019). The proportions of 15 BAs (with suffix ‘_p’) were calculated by dividing by total BA concentration. Additionally, 8 indices of BA metabolism were calculated (Chen et al., 2020): (1) Total BA (Total_BAs) = sum of all BA concentrations, (2) total primary BA (Total_primary_BAs) = sum of all primary BA concentrations, (3) total secondary BA (Total_secondary_BAs) = sum up of all secondary BA concentrations, (4) ratio of Secondary BAs to primary BAs ratio (Secondary_primary_ratio) = Total_primary_BAs/Total_secondary_BAs, (5) ratio of CA to CDCA concentrations (CA_CDCA_ratio) = (CA + TCA + GCA)/(CDCA + TCDCA + GCDCA), (6) ratio of unconjugated BA to conjugated BA concentrations = (CA + CDCA + DCA + LCA)/(TCA + GCA + TCDCA + GCDCA + TDCA + GDCA + TLCA + GLCA), (7) ratio of dehydroxylated CA to deconjugated CA concentrations (CA_dehydro_deconju_ratio) = (DCA + TDCA + GDCA)/(CA + TCA + GCA) and (8) ratio of taurine conjugated BA to glycine conjugated BA concentrations (Taurine_glycine_ratio) = (TCA + TCDCA + TDCA + TLCA)/(GCA + GCDCA + GDCA + GLCA).

### Metagenomic sequencing and quality control

Microbial DNA was isolated from fecal samples of LLD and 300-OB and sequenced as previously described (Kurilshikov et al., 2019; Zhernakova et al., 2016). We removed host genome□contaminated reads and low-quality reads from the raw metagenomic sequencing data using KneadData (version 0.7.4), Bowtie2 (version 2.3.4.3) (Langmead and Salzberg, 2012) and Trimmomatic (version 0.39) (Bolger et al., 2014). In brief, the data-cleaning procedure includes two main steps: (1) filtering out the human genome□contaminated reads by aligning raw reads to the human reference genome (GRCh37/hg19) and (2) removing adaptor sequences and low-quality reads using Trimmomatic with default settings (SLIDINGWINDOW:4:20 MINLEN:50).

### Taxonomic abundance

We generated the taxonomic relative abundance for both LLD and 300-OB samples from the cleaned metagenomic reads using MetaPhlAn3 with default parameters (Beghini et al., 2020).

### Detection of structural variations

Structural variants (SVs) are highly variable genomic segments within bacterial genomes that can be absent from the metagenomes of some individuals and present with variable abundance in other individuals. Based on the cleaned metagenomic reads, we detected the microbial SVs of all 1,437 samples from LLD and 300-OB using SGVFinder with default parameters. SGVFinder was devised and described by (Zeevi et al., 2019) and can detect two types of SV – deletion SVs (dSVs) and variable SVs (vSVs) – from metagenomic data. If the deletion percentage of the genomic segment across the population is < 25%, the standardized coverage will be calculated for this SV (vSV). If the deletion percentage is > 25% and < 75%, only the presence or absence status of this genomic segment will be kept (dSV). If the deletion percentage of a region is > 75%, the region is excluded from the analysis. The SV-calling procedure includes two major steps: (1) resolving ambiguous reads with multiple alignments according to the mapping quality and genomic coverage using the iterative coverage□based read assignment algorithm and reassigning the ambiguous reads to the most likely reference with high accuracy and (2) splitting the reference genomes into genomic bins and then examining the coverage of genomic bins across all samples to identify highly variable genomic segments and detect SVs. We used the reference database provided by SGVFinder, which is based on the proGenomes database (http://progenomes1.embl.de/) (Mende et al., 2017). In total, we detected 5,666 dSVs and 2,616 vSVs from 55 bacteria using default parameters. All bacterial species with SV calling were present in at least 5% of total samples.

### Functional annotation

The reference genome of *Blautia wexlerae* DSM 19850, *Coprococcus comes* ATCC 27758, *Eubacterium ventriosum* ATCC 27560, *Eubacterium hallii* DSM 3353 and *Eubacterium rectale* DSM 17629 were downloaded from progenome (http://progenomes1.embl.de/) (Mende et al., 2017) and annotated using the web-based genome annotation service provided by PATRIC (version 3.6.6, https://www.patricbrc.org/) (Wattam et al., 2017).

## QUANTIFICATION AND STATISTICAL ANALYSIS

All statistical tests were performed using R (version 4.0.1). Details of statistical tests are also provided in results and figure legends.

### Association analysis

Before association analysis, all continuous variables were standardized to follow a standard normal distribution (*N∼(0, 1)*) using empirical normal quantile transformation. Associations between SV and BAs were assessed in LLD and 300-OB using linear models with the following formula:

BA ∼ SV + Age + Sex + BMI + Reads number + Species relative abundance

The association between species relative abundance and BA parameters were assessed in LLD and 300-OB using linear model with following formula:

BA ∼ Species relative abundance + Age + Sex + BMI + Reads number

The association results of LLD and 300-OB were furtherly integrated using meta-analysis with a random-effect model, while the statistical heterogeneities were estimated with *I*^*2*^. To control the false discovery rate (FDR), Benjamina-Hochberg and Bonferroni P-value correction were performed using *p*.*adjust()* function in R. The association analysis and P-value correction were conducted for vSVs, dSVs and species relative abundance separately. The replicable significant SV□BA associations were confirmed with following four criteria: (1) P_LLD_ < 0.05, (2) P_300-OB_ < 0.05, (3) FDR_meta_ < 0.05 and (4) P_heterogeneity_ > 0.05.

The differences of BA parameters between SV-based clusters within species were tested using the Kruskal-Wallis rank-sum test. Empiric P values were estimated based on 999 permutations. For the analysis shown in **Figures S8C** and **S8D**, the Spearman correlation coefficient was calculated between the effect size in LLD and 300-OB. In **Figure S1A**, the mean value ± standard deviation is shown.

### Mediation analysis

The causal relationships between exposure factors, SVs and BAs were inferred by bidirectional mediation analysis with R package *mediation* (version 4.5.0). To reduce the number of tests, before mediation analysis we identified lifestyle□SV□BA groups in which all variables correlated with each other as candidate groups with a potential causal relationship. A candidate group had to meet the following criteria: (1) the association between the BA and SV is significant and replicable in both LLD and 300-OB, (2) the association between BA and lifestyle factor is significant (P < 0.05) and (3) the association between lifestyle factor and SV is significant (P < 0.05). We identified 1,338 candidate groups for vSVs and 175 candidate groups for dSVs. We then performed bidirectional mediation analysis on the candidate variable groups following the framework described in **Figure 6A**. For the vSV candidate groups, a linear model was used in each step of mediation analysis. For the dSV candidate groups, a logistic regression model was used when the response variable was a dSV. Finally, the P-values of indirect effects were corrected by FDR estimation.

### Distance calculation

We merged the vSV and dSV profiles and calculated Canberra distance between all samples based on the SV profile of each species respectively. We then standardized all matrices by dividing each matrix by its maximum distance value. To quantify the overall microbial genetic kinships between all individuals, we calculated the metagenome-wide genetic dissimilarities between all samples by calculating the distance of shared SVs. To quantify the overall compositional differences of the BA pool, we calculated the Canberra distance between all samples based on BA concentration profile and proportion profile. Distance matrices were computed using the *vegdist()* function from R package *vegan* (version 2.5-6).

### Unconstrained and constrained ordination analysis

We performed principal coordinates analysis (PCoA) on Canberra distance matrices of SV and BA profiles using *cmscale()* function from R package *vegan*. We estimated the proportion of BA pool variance explained by basic phenotypes (sex, age, and BMI) and cohort factor using permutational multivariate analysis of variance (PERMANOVA) with *adonis()* function from R package *vegan*. We estimated proportions of metagenome-wide SV-based genetic variance explained by age, sex, BMI and read count using PERMANOVA.

### Clustering analysis

Based on the genetic dissimilarity matrix of each species, we clustered the samples using the partitioning around medoid method and assigned samples to clusters with a given cluster number *k* (*k* ∈ [2, 10]). The best cluster numbers were determined by prediction strength (PS) (Tibshirani and Walther, 2012), with the highest number of clusters with a PS above 0.55 considered the best cluster number. If there was no PS value > 0.55, we assumed there was no obvious cluster (subspecies) within the corresponding species. The clustering results were then visualized using PCoA plot and t-distributed stochastic neighbor embedding (t-SNE) (Kobak and Berens, 2019).

